# Mapping galectin-3 ligands in tear fluid establishes spliceoform-dependent lacritin binding

**DOI:** 10.1101/2025.11.24.690288

**Authors:** Vincent Chang, Isaac Lian, Keira E. Mahoney, Jeff Romano, Ali Reza Afshari, Ryan J. Chen, Xiangjun Chen, Fredrik Fineide, Ayyad Zartasht Khan, Tor Paaske Utheim, Niclas G. Karlsson, Gordon W. Laurie, Stacy A. Malaker

**Affiliations:** Department of Chemistry, Yale University, New Haven CT, USA; Department of Life Sciences and Health, Faculty of Health Sciences, Oslo Metropolitan University, Oslo, Norway; Department of Cell Biology, University of Virginia, Charlottesville, Virginia, USA; Department of Ophthalmology, University of Virginia, Charlottesville, Virginia, USA; Department of Biomedical Engineering, University of Virginia, Charlottesville, Virginia, USA; Department of Plastic and Reconstructive Surgery, Oslo University Hospital, Oslo, Norway; Department of Ophthalmology, Vestre Viken Hospital Trust, Drammen, Norway; Department of Ophthalmology, Sørlandet Hospital Trust, Arendal, Norway; OsloMet, Oslo University Hospital, and the Dry Eye Clinic; Department of Medical Biochemistry, Oslo University Hospital, 0424 Oslo, Norway; Department of Ophthalmology, Østfold Hospital Trust, Moss, Norway

**Keywords:** galectin-3, lacritin, glycoproteomics, tear fluid, mass spectrometry, spliceoform

## Abstract

Galectin-3 (Gal-3) is a carbohydrate-binding protein which plays crucial roles in inflammation, immune response, cell migration, autophagy, and signaling. At the ocular surface, Gal-3 is also known to crosslink transmembrane mucins across the epithelial cell glycocalyx, forming lattice structures important for barrier function. However, the biological role of Gal-3 in circulating tear fluid remains largely unexplored. Similarly, whether Gal-3 engages extracellular glycoproteins in tears to affect downstream biological processes has yet to be investigated. As increased Gal-3 levels in tears are known to correlate with ocular pathologies such as dry eye disease (DED), we sought to elucidate the Gal-3 interactome in tear fluid and uncover biological insights into the function of Gal-3 beyond adhesion to the epithelial cell surface. Here, we combined ELISA, lectin affinity enrichment, mass spectrometry (MS)-based glycoproteomics, and lectin blotting to uncover Gal-3 interactors and their associated glycoepitopes. Overall, we report nearly 100 proteins enriched from tear fluid across 3 different patients, identifying proteins involved in immune response, inflammation, and antimicrobial activity. Most notably, we report lacritin as a novel ligand for Gal-3 and demonstrate that specific glycoforms of lacritin bearing core 2 O-glycans preferentially engage with Gal-3. Lastly, we show that the Gal-3-lacritin axis is spliceoform-specific and dependent on lacritin multimerization. Taken together, this study elucidates new ligands for Gal-3 in tear film and reveals mRNA splicing and multimerization as new biochemical mechanisms that fine-tune Gal-3 binding events.

## Introduction

Galectins comprise a family of β-galactoside-binding proteins that contain one or two conserved carbohydrate recognition domains (CRDs).^1^ Although β-galactose represents the minimal binding epitope, galectins display enhanced affinity toward N-acetyllactosamine (LacNAc), which can be found on core 2 O-glycans (GlcNAcβ1-6(Galβ1-3)GalNAcα-Ser/Thr) and complex or hybrid N-glycans.^2^ Galectins often engage glycoprotein ligands through LacNAc structures, and these interactions can promote cell adhesion, activate signaling pathways, and modulate the half-life of extracellular proteins.^1^ Given the biological significance of these binding events, the underlying mechanisms which regulate this process have been studied extensively.^3,4^ For instance, some mammalian galectins such as Gal-1,-2,-3, and −7 have the ability to dimerize, allowing for increased avidity towards carbohydrate antigens. Interestingly, Gal-3 is the only galectin which can also pentamerize, owing to its unique chimeric structure consisting of an N-terminal protein-binding domain in addition to its CRD.^5^ Here, the atypical structure of Gal-3 likely contributes to its involvement in a host of biological processes, including its ability to localize to the surface of microbes, activate immune cell receptors, and crosslink cell surface glycoconjugates.^6,7^ It is therefore unsurprising that Gal-3 is ubiquitously expressed across many different cell types and tissues and is often dysregulated in various diseases.^1,8^ As a result, over fifteen Gal-3 inhibitors have been investigated in clinical trials to date, with many targeting Gal-3 in cancer metastasis and other inflammatory diseases.^9^ However, despite extensive efforts in developing Gal-3 therapies, none have been explored thus far for ocular pathologies, largely due to a limited understanding of Gal-3 biology and its glycoprotein ligands in an ocular context.

Gal-3 is the only galectin known to be expressed at the ocular surface, and has been demonstrated to colocalize with MUC1 and MUC16 on the epithelial cell membrane.^10^ Furthermore, inhibition of ocular Gal-3 via galactose competition has been linked to increased membrane permeability and overall loss of barrier function. This supports a model in which multivalent Gal-3 crosslinks transmembrane mucins to form lattices that strengthen the mucosal barrier of the epithelial glycocalyx.^10,11^ As such, when barrier integrity is compromised during inflammation and pathogenic infiltration, a concomitant increase in Gal-3 levels in tear fluid is also observed.^12^ This is often seen in dry eye disease (DED), where tissue integrity and corneal maintenance is disrupted, resulting in chronic pain, impaired vision, and lasting corneal dysfunction.^13–15^ With the lack of molecular diagnostics and therapies for DED, further investigation into the therapeutic potential of ocular Gal-3 could be instrumental in guiding future treatment strategies.

While the concentration of Gal-3 in tear fluid is known to increase during inflammatory states, the biological roles and endogenous ligand(s) of Gal-3 beyond the epithelial cell surface remain largely unknown. Given that the tear film is rich in secreted glycoproteins derived from immune and epithelial cells, we hypothesized that Gal-3 binding events in the extracellular space were prevalent and could have downstream biological implications. Indeed, previous reports have demonstrated that Gal-3 binding to extracellular ligands such as fibronectin^16^, CD29^17^, CD66^18^, and α1β1 integrin^16^ are known to mediate cellular adhesion, T-cell apoptosis, and inflammation. With this in mind, we set out to uncover new Gal-3 ligands and their corresponding glycoepitopes in tear fluid. Here, we envisioned that elucidation of these binding interactions could inform therapeutic modalities targeted at Gal-3 in ocular diseases. Further, the functional role of site-specific glycans has yet to be investigated in the tear film, largely due to the high analytical complexity of this system. By leveraging recent advances in glycoproteomics^19–25^, we sought to reveal the biological significance of glycans through mapping the Gal-3 interactome.

Specifically, lacritin^26^ stands out within the tear fluid glycoproteome as an abundant glycoprotein (∼18 to 27 μM in tears)^27^ which plays key roles in tear production^28,29^, antimicrobial activity^30,31^, epithelial regeneration^32,33^, and protection against inflammatory stress^34^. Similar to Gal-3, lacritin expression levels are known to change significantly in DED^35^, coincident with increased inflammation and decreased lubrication.^27,13^ Recently, we reported the first in-depth view of the tear fluid glycoproteome, revealing site-specific O-glycosylation on lacritin with currently unknown function.^36^ Based on the glycosylation landscape of lacritin, we noted that core 2 O-glycoepitopes might serve as carbohydrate antigens for Gal-3 binding. In addition to elucidating lacritin glycoforms, we detected spliceoforms C and D of lacritin, representing the first observation of these isoforms by MS. Currently, it remains unclear if these splice variants might harbor distinct glycosylation profiles and whether they have different biological roles from the canonical isoform (A) of lacritin.^34^ Nonetheless, given that lacritin is ubiquitously expressed in tears and protects against inflammatory stress, we hypothesized that lacritin could be a novel ligand for Gal-3. Importantly, uncovering the major glycoepitopes on lacritin which engage Gal-3 could reveal a mechanistic basis for glycan-specific immunoregulatory events at the ocular surface, whereby dysregulation of this process may be concomitant with various diseases.

Several experimental methods have previously been explored for identifying potential Gal-3 ligands. For instance, Joeh et al. recently described a proximity-labeling approach to identify Gal-3 protein interactors in peripheral blood mononuclear cells.^37^ Alternatively, conventional immunoprecipitation and lectin-affinity enrichment methods have been widely used to capture Gal-3 ligands from complex samples.^38–42^ In either case, however, the site-specific glycoepitopes of the ligands often remain unknown without further experiments. To address these challenges, we leveraged a combination of lectin ELISA, MS-based affinity enrichment, glycoproteomics, and lectin blotting to define the Gal-3 interactome and site-specific glycoepitopes in tear fluid. Most notably, we establish lacritin as a ligand for Gal-3 and report specific lacritin glycosites which could engage Gal-3. Additionally, since lacritin is known to exhibit several splice variants^43^ (as described above) and exists as both a monomer and multimer^44^, we also assessed whether Gal-3 had different propensities for specific spliceoforms or multimers. Overall, this study serves to further our understanding of extracellular Gal-3 at the ocular surface by revealing key glycoproteins which engage Gal-3 and the underlying biochemical mechanisms which fine tune this interaction.

## Results and Discussion

### Investigation of Gal-3 binding in tear fluid with lectin ELISA and affinity-based enrichment

We first sought to quantify the extent of Gal-3 binding interactions (Gal-3 antigens shown in **Fig. 1A**) in tear fluid across a cohort of healthy and DED patients. To investigate this, we employed a recently developed fluorescent lectin assay (FLA) that enables quantitative measurement of lectin binding to biological fluids.^45^ Briefly, this assay leverages fluorescent europium (Eu) nanoparticles conjugated to a lectin which is applied to immobilized samples in a 96-well plate format (**Fig. 1B**). After a series of washes, a plate reader is used for fluorescent detection of lectin-bound samples. Here, we reasoned that this platform could be used to assess the presence of Gal-3 epitopes and their relative expression levels between healthy individuals (n=15) and DED patients with both low (n=15) and normal (n=15) tear fluid secretion. For our negative control, we immobilized an equal protein concentration of bovine serum albumin (BSA), which is not glycosylated. Overall, we observed a significant difference in Gal-3 binding between tear fluid glycoproteins and the BSA control across all three patient groups tested, indicating the presence of carbohydrate antigens that interact with Gal-3 (**Fig. 1B**). Interestingly, however, there was no significant difference in Gal-3 binding between the healthy and DED patient cohorts. To rationalize this observation, we note that previous proteomic studies showed global glycoprotein expression levels were similar between DED vs. healthy patients, as evidenced by the fact that less than 5 glycoproteins out of 1500 known tear proteins were dysregulated.^46,47^ On the other hand, the concentration of Gal-3 was significantly higher in DED.^12^ This likely suggests that Gal-3 binding events could be driven more by elevated Gal-3 expression levels as opposed to increased presentation of Gal-3 carbohydrate antigens in DED. Nonetheless, our data highlights significant Gal-3 binding in both healthy and DED patients.

**Figure 1.**
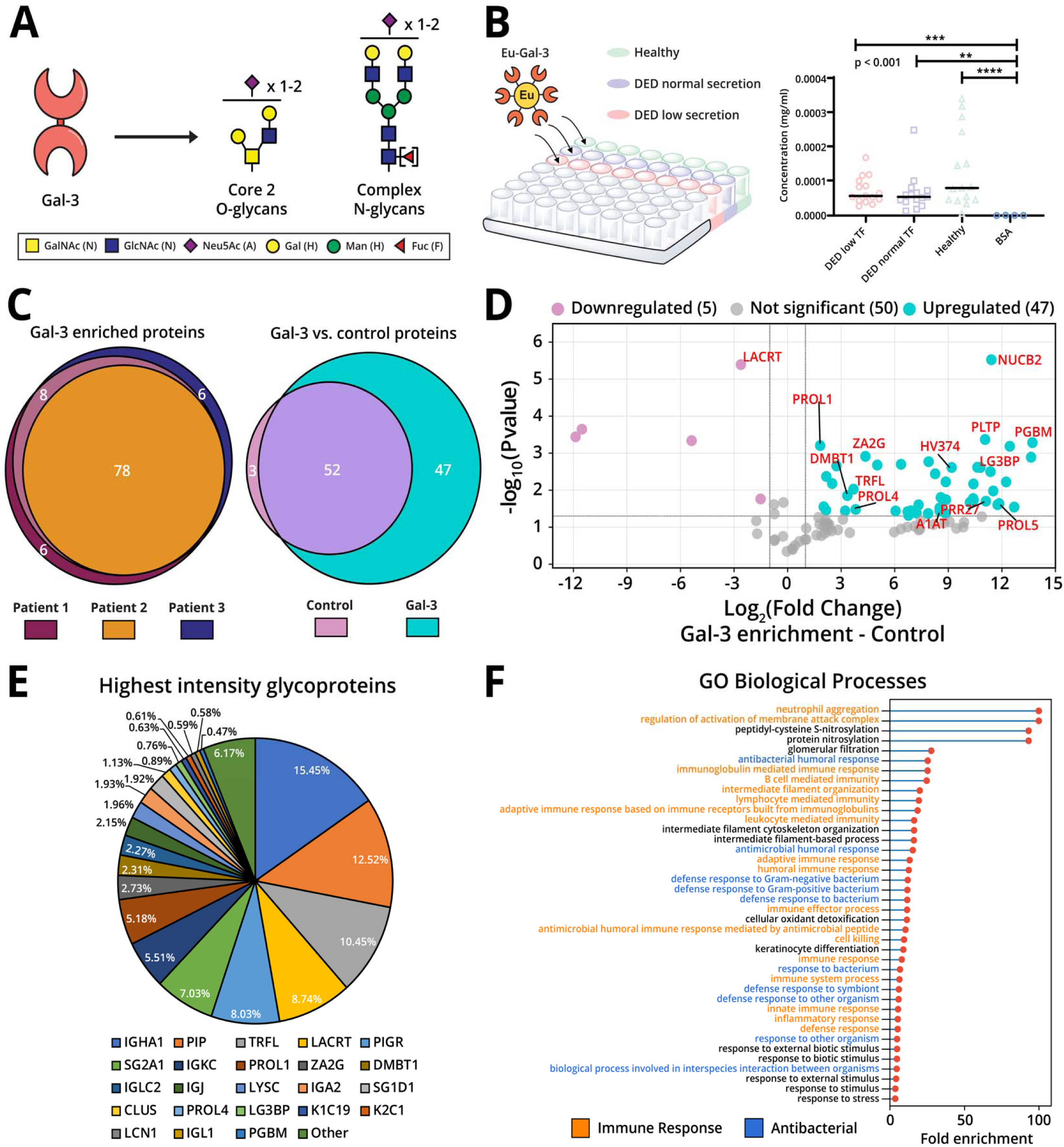
Gal-3 interactome analysis. **(A)** Gal-3 binds to core 2 O-glycans or complex type N-glycans. (**B**) Normalized Gal-3 concentration/binding to tear fluid glycoproteins and non-glycosylated BSA control. Statistical significance was calculated using a Kruskal-Wallis test. (**C**) Euler plot comparison of Gal-3-enriched proteins across 3 different patients (left) and compared to control (blocked) beads (right). Euler plot circles corresponding to patients 1,2, and 3 are colored in maroon, yellow, and blue respectively while control and Gal-3 enriched proteins are colored in pink and teal respectively. (**D**) Volcano plot displaying differentially enriched Gal-3 proteins compared to control in biological replicates. Cutoffs used for differentially expressed proteins are p<0.05 (as determined by a two tailed student’s t-test) and fold change >2. (**E**) Relative percent abundance analysis of the top 23 glycoproteins identified in all three Gal-3 enrichments. The full list of protein names can be found in **Table 1C**. (**F**) GO biological processes of identified proteins enriched by Gal-3 in at least two of three patients. Functions correlating to immune response and antibacterial activity are colored in orange and blue respectively.

After confirming the presence of carbohydrate antigens with FLA, we turned to proteomics to identify potential Gal-3 interactors. Here, we leveraged an affinity-based enrichment workflow where recombinantly expressed Gal-3 was immobilized onto a solid support. In brief, we incubated tear fluid from three donors with Gal-3-conjugated to N-hydroxy succinimide (NHS) agarose beads. After a series of washes, we eluted Gal-3 enriched proteins with an MS-compatible detergent sodium deoxycholate (SDC) before performing reduction and alkylation with dithiothreitol (DTT) and iodoacetamide (IAA).^48^ Finally, we added trypsin and the mucinase SmE (as performed in our previous study^36^) to generate peptides and glycopeptides for subsequent MS analysis (**Fig. S1**). Overall, we identified 99 tear proteins enriched by Gal-3 across the three donors, with 78 shared by all patients and 86 shared by at least two patients (**Fig. 1C, left, and Table 1A**). In parallel, we performed a control experiment where tear fluid from the three patients was rotated with blocked NHS solid support and then subjected to identical downstream processing. Notably, we identified 47 unique proteins and 52 shared proteins in the Gal-3 enriched samples when compared to the control (**Fig. 1C, right, and Table 1B, C**). To visualize statistically significant enriched proteins (p<0.05 and Fold change >2), we constructed a volcano plot (**Fig. 1D, Table 1D**). Here we highlighted several known tear fluid glycoproteins spanning immunoglobulins, proline-rich proteins (PRPs), lactoferrin (TRFL), and densely O-glycosylated proteins such as deleted in malignant brain tumors 1 (DMBT1), lacritin (LACRT), and heparan sulfate proteoglycan core protein (PGBM). To better understand the shared functions of these glycoproteins, we performed gene ontology (GO) analysis and discovered that enriched glycoproteins were predominantly involved in immune response, inflammation, and antibacterial activity (**Fig. 1F, table 1E**). Based on the processes listed in **Fig. 1F**, it is plausible that Gal-3 circulating in tears may serve a role in interacting with glycoproteins derived from neutrophils and immune cells near sites of ocular inflammation. Elevated levels of Gal-3 proximal to these sites might act to increase the local concentration of immunoglobulins and antimicrobial proteins in response to pathogens and stress during infection and disease. This would corroborate previous observations of higher Gal-3 levels in tears of DED patients^12^, given that DED is known to be inflammatory and can be driven by bacterial infections.^13,14,49^

Finally, we asked whether lacritin might serve as a ligand for Gal-3. Interestingly, we noticed lacritin was lower in abundance relative to the control according to the volcano plot. Nonetheless, lacritin still constituted 8.74% of the total protein intensity in Gal-3 enriched samples (**Fig. 1E**) and was the 4^th^ most abundant protein, suggesting significant binding to Gal-3. To account for the difference in overall lacritin intensity between Gal-3 enriched samples and the control, we hypothesized that only a subpopulation of lacritin glycoforms displaying core 2 O-glycans might serve as Gal-3 ligands.

### Glycoproteomic analysis reveals site-specific glycoepitopes enriched by Gal-3

To build on our initial proteomic analysis, we wanted to dive deeper into our dataset to investigate the site-specific glycosylation signatures enriched by Gal-3. First, we validated that the total glycosylation intensity and chromatographic profiles across the three biological replicates were similar. During higher energy collisional dissociation (HCD), oxonium ions are generated from glycans which fragment from a given glycopeptide. This can then serve as a general proxy for the extent of glycosylation in a sample.^50,51^ Here, we profiled HexNAc and Neu5Ac oxonium ions by extracting MS2 ion chromatograms corresponding to m/z values 138.055,144.065, 204.087, 274.092, and 292.103 (**Fig. S2A and S2B**). Overall, we observed similar chromatographic traces and ion current intensities (**Fig. S2C**), which validated that our biological replicates had comparable glycosylation profiles. Next, we analyzed different N-glycan structures that were captured by Gal-3 enrichment, anticipating Gal-3 would primarily bind N-glycans containing LacNAc. As expected, unique N-glycopeptides (i.e., glycopeptides with distinct sequence and/or glycan structure) markedly increased after enrichment, with complex type increasing roughly seven-fold (7 to 50) and hybrid type rising from 0 to 10. (**Fig. 2A, table 1F**). These N-glycopeptides primarily derived from immunoglobulins, corroborating our previous observations of enriched proteins involved in immune response and inflammation.

**Figure 2.**
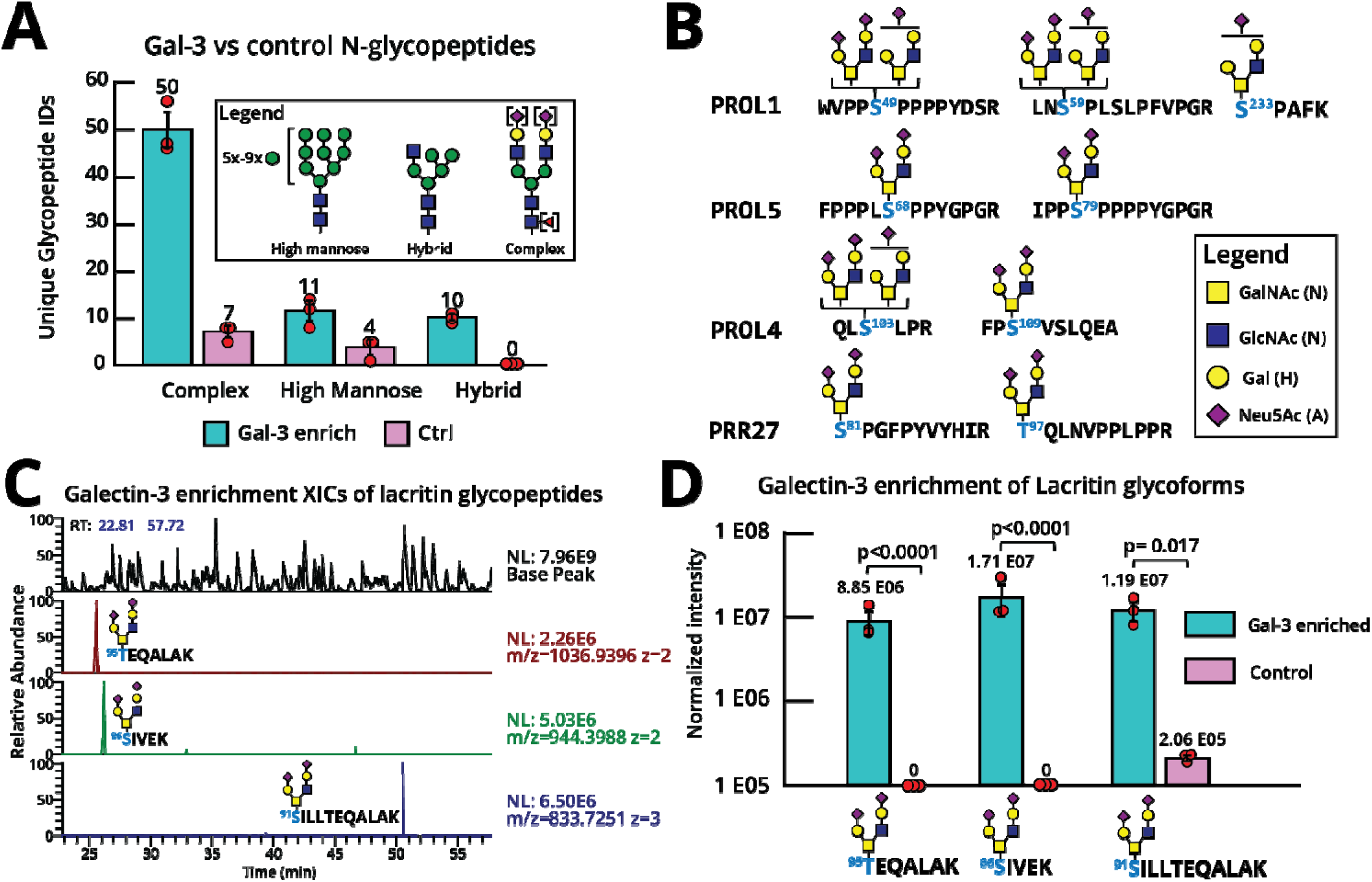
In-depth glycoproteomic analysis. **(A)** Full sequence of the canonical isoform of lacritin. The signal sequence is underlined and shaded in gray, localized O-glycosites are colored in blue, and the N-glycosite is colored in orange. Regions important for cell adhesion (orange), syndecan binding (blue), and representing the mucin domain (purple) are also shown. **(B)** Site-specific quantification of lacritin at each glycosite (left). Relative O-glycan abundances across all O-glycosites. Quantification was performed using label free quantitation (LFQ) and area under the curve (AUC) values from extracted ion chromatograms (XICs) of all identified glycopeptides.

In performing a similar analysis for O-glycopeptides bearing LacNAc, we note that none were identified in control samples, whereas Gal-3 enriched samples contained numerous PRPs (PROL1, PROL4, PROL5, PRR27) bearing core 2 O-glycans (**Fig. 2B, table 1G**). Historically, PRPs in saliva and tear fluid have been proposed to serve as a source of antimicrobial peptides owing to their high proline content.^52^ Though the role of glycans on proline rich proteins is not known, it is plausible that their glycans may affect proteolytic cleavage sites to regulate the length of antimicrobial peptides. Based on our results, it is reasonable to infer that core 2 O-glycans on PRPs can serve as ligands for Gal-3. Interestingly, Gal-3 is known to localize to the surface of microbes^23,24^ and is also highly expressed by immune cells and macrophages.^1^ It is therefore possible that interactions between Gal-3 and PRP glycoepitopes affect the local concentration of PRPs or alter their extracellular half-life, as has previously been shown for other extracellular glycoproteins.^1,8^ In addition to PRPs, we identified lacritin glycoforms bearing di-sialylated core 2 O-glycans across all three Gal-3-enriched patient samples (**Fig. 2C**). To validate the lacritin glycopeptides **S**[H2N2A2]IVEK, **T**[H2N2A2]EQALAK, and **S**[H2N2A2]ILLTEQALAK, we extracted ion chromatograms (XICs) for each precursor and further inspected MS2 fragment ions to confirm their sequence assignment and glycan localization (see **Fig. S3** for an example of an annotated MS2 spectrum). As noted previously in **Fig. 2B**, the total lacritin intensity decreased after Gal-3 enrichment. However, we observed that lacritin glycoforms bearing core 2 O-glycans were detected at significantly higher intensities relative to the control, as quantified by taking the area under the curve (AUC) intensity of each glycopeptide across all three samples (**Fig. 3D**). This observation rationalizes the decrease in lacritin intensity post-enrichment, suggesting that specific lacritin glycoforms harboring core 2 O-glycans preferentially bind Gal-3.

**Figure 3.**
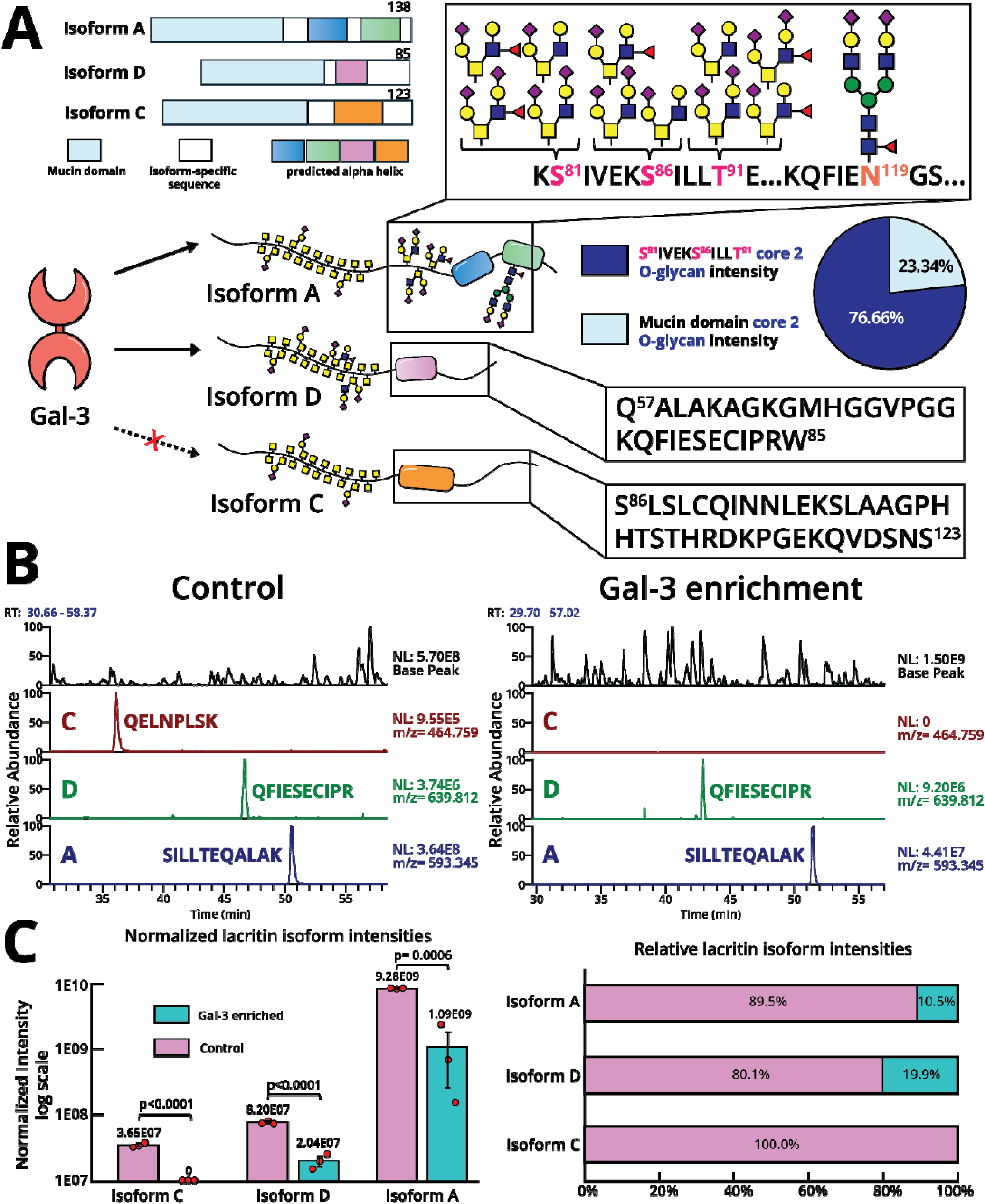
Gal-3 binds to isoforms A and D of lacritin. (**A**) Cartoon representation of lacritin splice variants and their spliceoform-specific regions (left). Representative glycan structures which were previously identified on Isoform A contain Gal-3 glycoepitopes, while isoform C and D were not identified to contain core 2 O-glycans in their C-terminal half (right). Relative core 2 O-glycan intensities on SIVEKSILLT and the mucin domain were also quantified by taking AUC intensities (middle, right). (**B**) XICs of spliceoform-specific tryptic peptides in a control enrichment (left) and a Gal-3 enriched (right) sample. (**C**) Bar graph representation of relative isoform abundances in the control and Gal-3 enriched samples. AUC intensities of each isoform specific peptide were normalized against a tryptic lacritin peptide (QFIENGSEFAQK) present across all three biological replicates. Error bars represent standard error of mean and statistical significance is calculated with a two-tailed student’s t-test.

### Gal-3 engages specific lacritin spliceoforms

As mentioned previously, lacritin undergoes alternative splicing to generate isoforms C and D in addition to the canonical sequence (isoform A). In our prior study, we reported MS-based detection of isoform-specific tryptic peptide sequences corresponding to isoforms A (SILLTEQALAK), C (QELNPLSK), and D (QFIESCIPR) of lacritin.^36^ We further hypothesized that different lacritin splice variants might harbor distinct glycosylation profiles and therefore exhibit unique biological roles or engage different protein ligands. With this in mind, we asked whether Gal-3 preferentially engaged specific lacritin splice variants. As shown in **Fig. 2D**, we observed that the enriched core 2 O-glycopeptides were unique to a spliceoform-specific exon of isoform A (**Fig. 3A**). Furthermore, Ser86, Ser91, and Thr95 contained within this exon had a significantly higher relative abundance of core 2 O-glycans (76.66% of the core 2 O-glycan intensity of lacritin) compared to the densely O-glycosylated N-terminal half (23.34%), indicating that Gal-3 primarily engages glycans across these 3 residues (**Fig. 3A**). Surprisingly, we did not detect any isoform C-specific tryptic peptides across all three patients after Gal-3 enrichment (**Fig. 3B, Fig. S4**). Based on the protein sequence of isoform C, 6 Ser residues in the C-terminal region could serve as potential glycosites (**Fig. 3A**). However, in the enriched samples, we were unable to detect any glycopeptides corresponding to these glycosites and therefore speculate that isoform C is unlikely to bear Gal-3 glycoepitopes. Conversely, isoform A and D were observed to bind Gal-3 in all replicates and patients (**Fig 3B, Fig. S4**). Protein sequence analysis of isoform D revealed a lack of C-terminal glycosites, suggesting that its densely O-glycosylated N-terminal mucin domain is likely to display the core 2 O-glycoepitopes for Gal-3 binding (**Fig. 3A**). Alternatively, protein-protein crosslinking between isoform D and A to form heterodimers may also explain Gal-3 enrichment of this isoform. To estimate the relative lacritin populations of each isoform which can engage Gal-3, we performed AUC quantification and observed approximately 10.5% of isoform A, 19.9% of isoform D, and 0% of isoform C likely contains core 2 O-glycans (**Fig. 3C**) when compared to unenriched controls.

Discovery of lacritin as a spliceoform-specific ligand for Gal-3 has important implications for uncovering the roles of different lacritin isoforms in tear fluid. To date, the biological function and significance of isoforms C and D remain unknown,^27^ though the lack of Gal-3 binding to lacritin-C suggests that isoforms A and D may exhibit distinct biological functions. While lacritin-A is well characterized and is known to participate in mitogenesis, antimicrobial activity, and tear production, the existence of isoform D at the protein level was only recently reported by our group, having previously thought to not be translated.^36^ Given that lacritin-D serves as a ligand for Gal-3, it may also serve as a source of antimicrobial peptides or engage other protein receptors at the immune interface.

More broadly, mRNA splicing as a regulatory mechanism for fine-tuning the display of glycoepitopes has been well documented across other glycoproteins. For instance, CD34^54^, CD44^55,56^, CD45^57^, MUC1^58^, and MUC4^59^ are all known to undergo alternative splicing to generate spliceoforms with different glycosylation profiles due to the loss of an exon containing glycosites or a variation in tandem repeat length. To the best of our knowledge, however, lacritin is the first example of a glycoprotein whereby different splice variants harbor distinct binding propensities to a galectin. Moving forward, we envision other biological examples of this phenomenon will be discovered and will continue to unravel the significance of isoform-specific glycan-binding events.

### Lectin blotting reveals lacritin multimers and lactoferrin as a ligand for Gal-3

Previously, the Laurie group showed that lacritin monomers could undergo multimerization through transglutaminase-mediated crosslinking to form dimers, trimers, and larger polymers.^44^ This process has significant implications for the biological activity of lacritin, which is mediated by its ability to bind syndecan-1 (the only known ligand for lacritin).^60–62^ In this study, it was revealed that only monomeric lacritin bound syndecan-1, suggesting that the multimeric state of lacritin could regulate its affinity toward a ligand. These observations led us to investigate the possibility that both lacritin multimers and monomers might serve as Gal-3 ligands and potentially exhibit distinct binding affinities.

However, while we were able to establish lacritin as a ligand by bottom-up proteomics using the methods described above, it was not possible to distinguish whether multimeric or monomeric lacritin had engaged Gal-3. To address this, we first immunoprecipitated all forms of lacritin from 100 µL of pooled tear fluid with a panel of lacritin antibodies (**Fig. 4A**). After immunopurification with protein A/G spin columns, lacritin was loaded onto an SDS-PAGE gel before lectin blot analysis with His-tagged Gal-3 and a fluorescent (IR-800) anti-His antibody. Interestingly, while we observed Gal-3 binding to both monomeric and multimeric lacritin, multimers seemed to exhibit higher binding as shown by a more intense band on the Gal-3 lectin blot (**Fig. 4B**). We then performed quantitative densitometry on three replicates of pooled tear fluid. From this analysis, we estimated that dimeric and trimeric lacritin have a ∼3.5 fold and ∼2 fold higher affinity for Gal-3, respectively, compared to the monomeric form. Based on this result, it is anticipated that lacritin multimers may harbor different glycosylation profiles from monomers. Another explanation for the difference in Gal-3 binding could be due to avidity, whereby multimers contain more total glycosites than monomers and are therefore capable of displaying more core 2 glycoepitopes to engage Gal-3. We note, however, that this could not account for dimeric lacritin exhibiting a higher Gal-3 affinity than trimeric lacritin. Given that bottom-up glycoproteomics cannot distinguish between monomer vs. multimer glycoepitopes with current methods, future efforts will be necessary to isolate these individual species to investigate their glycosylation differences separately. Uncovering multimer and monomer-specific glycan profiles could account for the unique preferences in Gal-3 binding observed in this study and provide insight into differences in the structure and function between single copies and crosslinked versions of lacritin.

**Figure 4.**
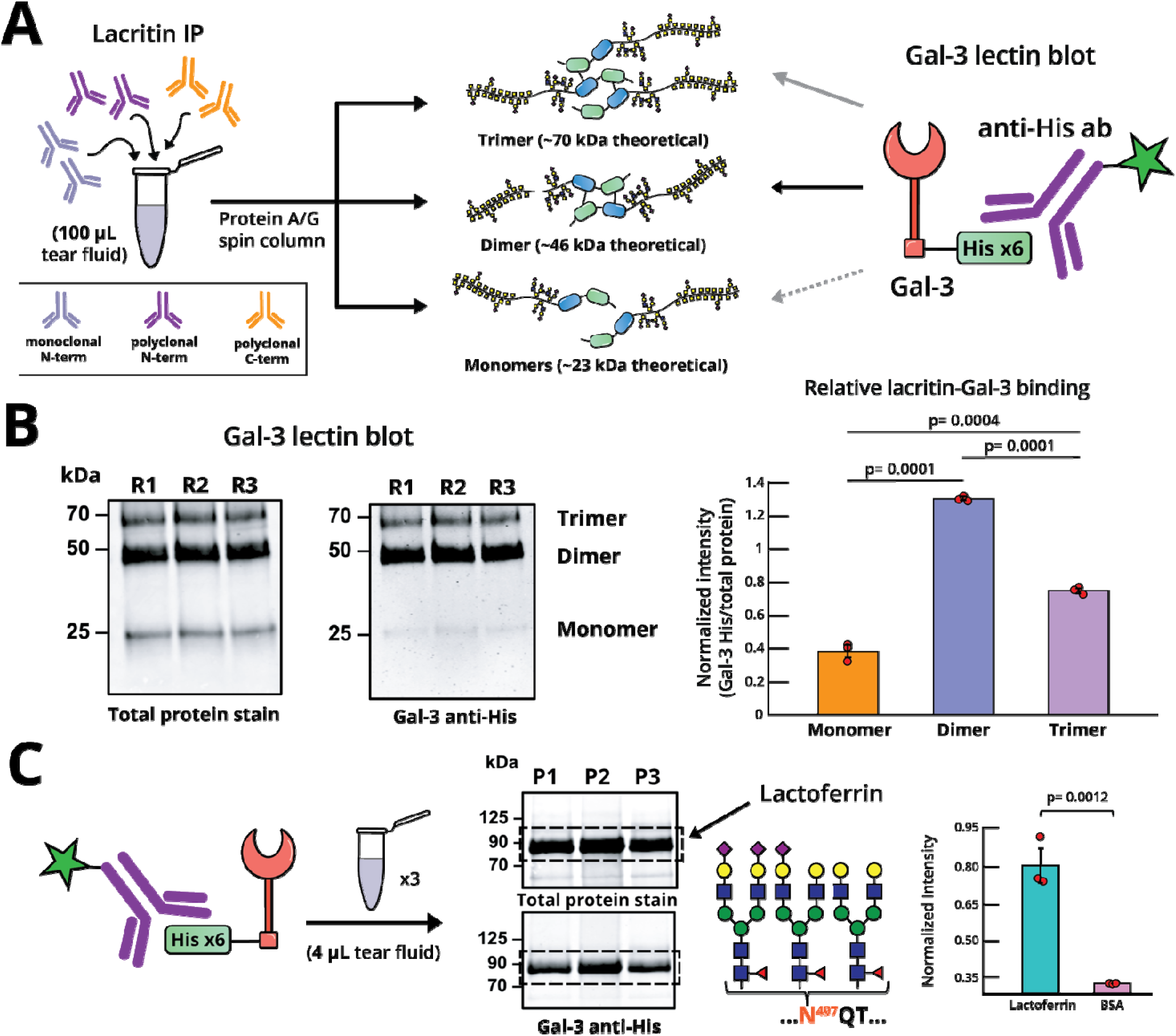
Gal-3 blotting of lacritin and lactoferrin for ligand validation. (**A**) Immunoprecipitation of lacritin from pooled patient tear fluid using a panel of three lacritin antibodies (left). Immunopurified lacritin exists as monomers, dimers, or trimers (middle) which can then be probed for Gal-3 binding by a Gal-3 lectin blot (right). (**B**) Replicate analysis of pooled patient tears (denoted R1, R2, and R3 as replicates 1, 2, and 3). REVERT total protein stain shows lacritin bands migrating at ∼70 kDa, ∼46 kDa, and ∼23 kDa corresponding to the trimer, dimer, and monomer respectively. Gal-3 anti-His shows staining of each band, with dimeric and trimeric lacritin having higher band intensity when compared to monomeric lacritin (left). Densitometry analysis of Gal-3 binding to lacritin, where the normalized intensity is taken as the band intensity in Gal-3 anti-His divided by the total protein stain intensity of the same band. (**C**) Gal-3 blot on tear fluid of three individual patient donors (denoted as P1, P2, and P3 as patients 1, 2, and 3) reveals lactoferrin (at ∼90 kDa) as a ligand for Gal-3 (middle). Densitometry analysis (right) showing normalized intensity difference between lactoferrin and nonglycosylated BSA control, quantified in the same way as (**B**).

In addition to lacritin, lactoferrin is a highly expressed glycoprotein with known antibacterial activity, anti-inflammatory properties, and the ability to promote cell proliferation.^63,64^ Lactoferrin contains three N-glycosites (Asn156, Asn497, and Asn642), with Asn497 being the most frequently (99%) occupied.^65,66^ Previously, we reported both complex (96% relative abundance) and high mannose (4% relative abundance) N-glycan structures on Asn497.^36^ As the role of lactoferrin N-glycans in tear fluid remains unknown, we hypothesized that the presence of complex N-glycans on Asn497 might engage Gal-3. From our initial Gal-3 affinity enrichment, we noticed lactoferrin was significantly enriched in our volcano plot (**Fig. 1B**) and constituted a significant portion of the total protein signal post-enrichment (3^rd^ most abundant protein; **Fig. 1C**). Furthermore, due to the high abundance of lactoferrin in tears, numerous studies have characterized a distinct high-intensity band in SDS-PAGE at ∼90 kDa as corresponding to lactoferrin in non-reducing conditions.^67–69^ These observations prompted us to analyze tear fluid from three donors where we similarly observed a discrete and intense band corresponding to lactoferrin in the total protein stain (**Fig. 4C**). We then confirmed Gal-3 staining of lactoferrin bands via lectin blotting. To ensure that this interaction was not the result of non-specific binding, we compared the normalized intensity of lactoferrin binding to a non-glycosylated BSA control and observed significantly higher intensities (**Fig. 4C**). Validation of this binding interaction has implications for a new potential function of lactoferrin N-glycosylation through modulation of Gal-3 binding in tears. As with lacritin, this binding interaction may affect the local concentration of lactoferrin or alter the proteolytic processing of this protein into antimicrobial peptides. Interestingly, lactoferrin and Gal-3 are co-expressed in many other biofluids and tissues across the body.^64,70,71^

## Conclusion

In this study, we hypothesized the existence of extracellular Gal-3 binding events beyond the epithelial cell surface in tears. To explore this, we first performed FLA to validate the presence of Gal-3 carbohydrate antigens in tear fluid across a cohort of healthy individuals and DED patients. We then leveraged MS-based affinity enrichment to identify Gal-3 interactors, revealing numerous proteins involved in immune response and antimicrobial activity. Within the tear fluid glycoproteome, lacritin emerged as an abundantly expressed glycoprotein with key roles in immune response, tear secretion, and antimicrobial activity. Given that lacritin was previously shown to display core 2 O-glycans and the fact that both Gal-3 and lacritin expression levels are dysregulated in DED, we investigated whether Gal-3 could be a novel ligand for lacritin. Through a combination of glycoproteomics, immunoprecipitation, and lectin blotting, we discovered that the Gal-3-lacritin axis is glycoform- and spliceoform-specific. Surprisingly, it is also dependent on the multimeric state of lacritin. Future studies to characterize the glycosylation landscape of multimers separately from monomers will be crucial to shed light on potential functional and structural differences which are driven by site-specific O-glycans.

The elucidation of new Gal-3 ligands and their corresponding glycoepitopes in tear fluid helps to inform strategies for selective therapies targeted at Gal-3 in the ocular space. Beyond these therapeutic implications, we offer new biochemical insights into mechanisms that can dictate glycan-protein interactions. Most notably, we highlight alternative protein splicing and multimerization of lacritin as two processes which fine-tune its binding affinity towards Gal-3. Moving forward, it will be crucial to account for these events when characterizing new ligands of Gal-3 and other galectins in separate biological contexts.

Taken together, this study represents the first investigation into the functional significance of site-specific glycans in tear fluid. As the tear-fluid glycoproteome has only recently been defined by glycoproteomics, further studies will be crucial to uncover the rich glycobiology underlying this system. Our results lay the critical groundwork for future investigation into the downstream biological consequences of Gal-3 binding. Moreover, the functional role of lacritin isoforms and multimers still remains unknown, though we now know these lacritin populations exhibit different binding affinities towards Gal-3.

## Materials and Methods

### Tear fluid sample collection for fluorescence lectin assay

Tear fluid from DED and healthy patients were collected via Schirmer strips. DED patients with low tear fluid production were defined as symptomatic individuals (Dry Eye Questionnaire 5 [DEQ-5] > 7 or Ocular Surface Disease Index [OSDI] > 12) and a fluorescein tear-film break-up time (TBUT) < 10 seconds and a Schirmer test (ST) < 10 mm. DED patients with normal tear fluid production were defined as symptomatic individuals (DEQ-5 > 7 or OSDI > 12) and a TBUT < 10 seconds and a Schirmer test (ST) > 10 mm. Healthy controls were defined as asymptomatic participants (DEQ-5 < 7 and/or OSDI < 12) and a TBUT > 10 seconds. All clinical measurements and tear samples were obtained from the left eyes (OS). A total of 45 participants were included: DED with low tear fluid production (n = 15; 3 males; age range = 23–78 years), DED with normal tear fluid production (n = 15; 4 males; age range = 23–77 years), and healthy controls (n = 15; 11 males; age range = 24–66 years). Participants were recruited through the Norwegian Dry Eye Clinic and various Specsavers clinics in Norway. All participants provided informed consent. The study (including the analyses of the biological samples) was approved by the Regional Committee for Medical and Health Research Ethics (REK) in South-Eastern Norway (approval numbers 427997 and 269845).

### Fluorescence lectin assay

Proteins from Schirmer strips were extracted in 200 µl urea (7M), thiourea (2 M) and the protein concentrations were determined by Bradford assay. Proteins (37.5 ng in 25 µl) were statically coated in each well on Nunc-Immunomaxisorp plates (Thermo Fisher Scientific) (Overnight @ 4°C). After blocking with 1% BSA (100 µl/well, 45 min RT) lectin binding was detected using Gal-3 conjugated Europium nanoparticles (50 µl) as previously described.^45^

### Tear fluid samples for MS analysis, lacritin immunoprecipitation, and lectin blot

Tear fluid for MS analysis, lacritin IP, and lectin blot was purchased from Innovative Research (obtained via microcapillary collection) and stored at −20°C until further processing. 1 pooled sample was used for lacritin IP and 3 individual donors (56669-ND0557-CF35, 56668-ND0350-CF42, and 56667-ND0143-CF43) were used for MS and lectin blot analysis.

### Mass spectrometry sample preparation

#### Preparation of Galectin-3 beads

The pET28 plasmids for His-tagged SmE and Gal-3 was kindly provided by the Bertozzi laboratory. NHS-activated beads were first washed 3x with 1mL of PBS. Next, Gal-3 (1 mg/mL in 1mL PBS) was conjugated to NHS-activated beads (500 µL bead slurry) overnight with agitation at 4 °C. After removing the supernatant the following day, free NHS-esters were capped by adding 100 mM Tris, pH 7.4, to the bead slurry for 20 min at 4 °C. BCA assays were performed after the reactions in order to determine the sufficient binding efficiency. After capping, beads were washed 3x with a high-salt buffer (20 mM Tris and 500 mM NaCl) followed by a buffer without salt (20 mM Tris).

#### Galectin-3 affinity-based enrichment

Tears (diluted to 500 µL with PBS with a final concentration of 0.5 mg/mL) were loaded onto Gal-3-conjugated NHS beads (or NHS beads blocked with 100mM tris for control enrichments), washed 3 times with 20 mM Tris and 500 mM NaCl, and eluted with 0.5% SDC. Following elution, the mucin-enriched tear fluid was subjected to reduction (2mM DTT at 65 °C for 1 hour), alkylation (5mM IAA at 23 °C in the dark for 15 minutes), and proteolytic digestion with mucinase SmE (E:S ratio of 1:20 overnight at 37 °C) and trypsin (E:S ratio of 1:50 for 6 hours at 37 °C). All reactions were quenched by adding 1 µL of formic acid and diluted to a volume of 200 µL prior to desalting. Addition of formic acid also caused SDC to precipitate out of solution, and the resulting supernatant was transferred to a new tube before desalting. Desalting was performed using 10 mg Strata-X 33 µm polymeric reversed phase SPE columns (Phenomenex). Each column was activated using 500 µL of acetonitrile (ACN) (Honeywell) followed by of 500 µL of 0.1% formic acid, 500 µL of 0.1% formic acid in 40% ACN, and equilibration with two additions of 500 µL of 0.1% formic acid. After equilibration, the samples were added to the column and rinsed twice with 200 µL of 0.1% formic acid. The columns were transferred to a 1.5 mL tube for elution by two additions of 150 µL of 0.1% formic acid in 40% ACN. The eluent was then dried using a vacuum concentrator (LabConco) prior to reconstitution in 10 µL of 0.1% formic acid. The resultant peptides were then injected onto a Dionex Ultimate3000 coupled to a Thermo Orbitrap Eclipse Tribrid mass spectrometer. We employed a higher-energy collision dissociation product-dependent electron transfer dissociation (HCD-pd-ETD) method; in some cases, we used supplemental activation in ETD (EThcD). The files were searched using Byonic, followed by manual data curation.

### Unenriched tear fluid processing

Tear fluid proteins from each patient were diluted to a final concentration of 0.2 mg/mL in 100 µL of PBS. DTT was then added to a concentration of 2 mM and reacted at 65 °C for 1 hour followed by alkylation in 5 mM IAA for 15 min in the dark at RT. Subsequently, digestion with trypsin was done at a 1:50 E:S ratio for 6 hours at 37 °C. All reactions were quenched by adding 1 µL of formic acid and diluted to a volume of 200 µL prior to desalting. Desalting and sample injection into the mass spectrometer was performed as previously described in the GlycoFASP methods section.

### Mass spectrometry data acquisition

Samples were analyzed by online nanoflow liquid chromatography-tandem mass spectrometry using an Orbitrap Eclipse Tribrid mass spectrometer (Thermo Fisher Scientific) coupled to a Dionex UltiMate 3000 HPLC (Thermo Fisher Scientific). For each analysis, 4 µL was injected onto an Acclaim PepMap 100 column packed with 2 cm of 5 µm C18 material (Thermo Fisher, 164564) using 0.1% formic acid in water (solvent A). Peptides were then separated on a 15 cm PepMap RSLC EASY-Spray C18 column packed with 2 µm C18 material (Thermo Fisher, ES904) using a gradient from 0-35% solvent B (0.1% formic acid with 80% acetonitrile) in 60 min. Full scan MS1 spectra were collected at a resolution of 60,000, an automatic gain control target of 3e5, and a mass range from *m/z* 300 to 1500. Dynamic exclusion was enabled with a repeat count of 2, repeat duration of 7 s, and exclusion duration of 7 s. Only charge states 2 to 6 were selected for fragmentation. MS2s were generated at top speed for 3 seconds. Higher-energy collisional dissociation (HCD) was performed on all selected precursor masses with the following parameters: isolation window of 2 m/z, 29% normalized collision energy, orbitrap detection (resolution of 7,500), maximum inject time of 50 ms, and a standard automatic gain control target. An additional electron transfer dissociation (ETD) fragmentation of the same precursor was triggered if 1) the precursor mass was between m*/z* 300 to 1500 and 2) 3 of 8 HexNAc or NeuAc fingerprint ions (126.055, 138.055, 144.07, 168.065, 186.076, 204.086, 274.092, and 292.103) were present at *m/z* ± 0.1 and greater than 5% relative intensity. Two files were collected for each sample: the first collected an ETD scan with supplemental energy (EThcD) while the second method collected a scan without supplemental energy. Both used charge-calibrated ETD reaction times, 100 ms maximum injection time, and standard injection targets. EThcD parameters were as follows: Orbitrap detection (resolution 7,500), calibrated charge-dependent ETD times, 15% nCE for HCD, maximum inject time of 150 ms, and a standard precursor injection target. For the second file, dependent scans were only triggered for precursors below *m/z* 1000, and data were collected in the ion trap using a normal scan rate.

### Mass spectrometry data analysis

Raw files were searched using Byonic (version 4.5.2, Protein Metrics, Inc.) against the UniProtKB/Swiss-Prot *Homo sapiens* proteome (Query: proteome:up000005640 AND reviewed:true) and a curated mucin database which was generated from a previous study. Briefly, the mucin database consists of nearly 350 proteins from Uniprot’s annotated human proteome predicted to bear the dense O-glycosylation characteristic of mucin domains. For all samples, we used the default O-glycan database containing 9 common structures. Raw files were first searched against the human proteome and then the curated mucin database. In both cases, files were searched with semi-specific cleavage N-terminal to Ser and Thr and six allowed missed cleavages. Samples treated with trypsin were searched with the same parameters but also allowed cleavage C-terminal to Arg or Lys. Mass tolerance was set to 10 ppm for MS1’s and 20 ppm for MS2’s. Met oxidation was set as a variable modification and carbamidomethyl Cys was set as a fixed modification. From the Byonic search results, glycopeptides were filtered to a score of >200 and a logprob of >2. From the remaining list of glycopeptides, the extracted ion chromatograms, full mass spectra (MS1s), and fragmentation spectra (MS2s) were investigated in XCalibur QualBrowser (Thermo) to generate a list of true-positive glycopeptides, as reported in **Table 1**. Each reported glycopeptide listed in **Table 1** was manually validated from the filtered list of Byonic’s reported peptides (score>200 and logprob >2) according to the following steps: The MS1 was first used to confirm the precursor mass and chosen isotope was correct. This also allowed us to identify any co-isolated species that could interfere with the MS2s and/or explain unassigned peaks. The HCD and EThcD fragmentation spectra were then investigated to identify sufficient coverage to make a sequence assignment. When possible, multiple MS2 scans were averaged to obtain a stronger spectrum. For HCD, an initial glycopeptide identification was confirmed if the presence of the precursor mass without a glycan present (i.e., Y0), along with coverage of b and y ions without glycosylation. For longer peptides, we required the presence of Y0 and fragments that were expected to be abundant (e.g., N-terminally to Pro, C-terminally to Asp). When the peptide contained a Pro at the C-terminus, the b_n-1_ was considered sufficient. Further, when the sequence contained oxidized Met, the Met loss from the bare mass was considered as representative of the naked peptide mass. We then used electron-based fragmentation MS2 spectra for localization. Here, all plausible localizations were considered, regardless of search result output. We confirmed the presence of fragment ions in ETD or EThcD that were between potential glycosylation sites, if sufficient c/z ions were present then a glycan mass was considered localized. For glycopeptide manual validation, extracted ion chromatograms are evaluated at the MS1 level to determine the charge and m/z of the highest abundance precursor species. Mass spectrometry data files and raw search output can be found on PRIDE with identifier PXD064788.

### Lacritin immunoprecipitation

Antibodies for lacritin immunoprecipitation were monoclonal (anti-N-terminal 1F5), polyclonal N-terminal (‘anti-Pep Lac N-term’) and polyclonal C-terminal (‘ab C-term [‘anti-Pep Lac C-term’]).^43,72^ Immunoprecipitation was performed as a modification of a previous procedure.^34,43^ Briefly, the three lacritin antibodies were incubated overnight at 4 °C with 100 µL of tear fluid and brought to a final volume of 500 µL with PBS containing protease inhibitors. The following day, the mixture was added to protein A/G spin columns (Thermo: 89950) and lacritin was purified according to manufacturer’s protocol.

### Galectin-3 blot

4 µL of tear fluid or 30 µL of immunoprecipitated lacritin were run on a 4–12% Criterion XT BisTris gel (Bio-Rad, 3450123) in MES XT buffer (Bio-Rad, 1610789) at 180 V for 50 min, alongside Chameleon Duo Pre-stained Protein Ladder (LICORbio P/N: 928-60000). Proteins were then transferred onto a nitrocellulose filter paper with a Trans-Blot^®^ Turbo™ Transfer System (Bio-Rad, 1704150EDU) and stained with REVERT 700 total protein stain (LICORbio P/N: 926-11015) following manufacturer’s protocol. After total protein stain imaging, a 1.2% BSA blocking solution was allowed to rock with the blot paper for 1 hour at room temperature. Next, 5 µg/mL of His-tagged Gal-3 in 0.1% PBST was added to the blot paper and incubated at room temperature for 1 hour. Finally, anti-His antibody was added at a 1:1000 ratio for 1 hour after 4 washes with 0.1% PBST. After the secondary antibody was allowed to bind for 1 hour, the blot paper was washed twice with 0.1% PBST and once with PBS before imaging. All washes were done for 5-10 minutes and all protein blots were imaged on a LiCOR Odyssey instrument. Densitometry analysis was performed on ImageJ.

## Supporting information

This article contains supporting information.

## Data availability statement

All mass spectrometry data and search results acquired for this manuscript have been deposited on the PRIDE repository. Reviewers can access raw data with the following login information: Project accession: PXD071066, Token: iD7BJE9KjF3W Alternatively, reviewer can access the dataset by logging in to the PRIDE website using the following account details: Username: Password: f6AiaQz4kfK3

## Author contributions

V.C, and S.A.M. conceptualization; V.C., I.L. and A.R.A. data curation; V.C., I.L., A.R.A. and R.C. formal analysis; V.C. investigation; V.C., J.R., and K.E.M. methodology; V.C. validation; V.C. and I.L. visualization; V.C. and S.A.M. writing– original draft; V.C., R.C., A.R.A. J.R., X.C., F.F., A.Z.K., T.P.U., N.G.K., G.W.L. and S.A.M. writing–review & editing; X.C., F.F., A.Z.K., and T.P.U. patient screening and sample collection; S.A.M. supervision; N.G.K., G.W.L. and S.A.M. funding Acquisition; V.C. project administration; N.G.K., G.W.L. and S.A.M. resources.

## Funding and additional information

V.C. is supported by an NSF GRFP (DGE-2139841).S.A.M. is supported by CRI Lloyd J Old STAR Award and a NIGMS R35-GM147039. I.L. is supported by The Yale College First-Year Summer Research Fellowship in the Sciences and Engineering. We also gratefully acknowledge financial contribution to NGK from OsloMet’s seeding fund for internationalization 2024 (3000-116) and to TPU from Department of Medical Biochemistry, Oslo University Hospital, Oslo Norway. G.W.L. is supported by 2R01 EY026171 and R01 EY032956.

## Abbreviations

The abbreviations used are

ACN: Acetonitrile
ETD: Electron transfer dissociation
EThcD: Electron transfer/higher-energy collision dissociation
HCD: Higher-energy collisional dissociation
DED: Dry eye disease
PBS: Phosphate Buffered Saline

## Citations

(1) Johannes, L.; Jacob, R.; Leffler, H. Galectins at a Glance. J. Cell Sci. 2018, 131 (9), jcs208884. 10.1242/jcs.208884.

(2) Moure, M. J.; Gimeno, A.; Delgado, S.; Diercks, T.; Boons, G.-J.; Jiménez-Barbero, J.; Ardá, A. Selective 13C-Labels on Repeating Glycan Oligomers to Reveal Protein Binding Epitopes through NMR: Polylactosamine Binding to Galectins. Angew. Chem. Int. Ed. 2021, 60 (34), 18777–18782. 10.1002/anie.202106056.

(3) Earl, L. A.; Bi, S.; Baum, L. G. N- and O-Glycans Modulate Galectin-1 Binding, CD45 Signaling, and T Cell Death. J. Biol. Chem. 2010, 285 (4), 2232–2244. 10.1074/jbc.M109.066191.

(4) Cao, Z.; Said, N.; Amin, S.; Wu, H. K.; Bruce, A.; Garate, M.; Hsu, D. K.; Kuwabara, I.; Liu, F.-T.; Panjwani, N. Galectins-3 and −7, but Not Galectin-1, Play a Role in Re-Epithelialization of Wounds*. J. Biol. Chem. 2002, 277 (44), 42299–42305. 10.1074/jbc.M200981200.

(5) Tissue- and cell-specific localization of galectins, β-galactose-binding animal lectins, and their potential functions in health and disease | Anatomical Science International. https://link.springer.com/article/10.1007/s12565-016-0366-6 (accessed 2025-11-01).

(6) Feeley, E. M.; Pilla-Moffett, D. M.; Zwack, E. E.; Piro, A. S.; Finethy, R.; Kolb, J. P.; Martinez, J.; Brodsky, I. E.; Coers, J. Galectin-3 Directs Antimicrobial Guanylate Binding Proteins to Vacuoles Furnished with Bacterial Secretion Systems. Proc. Natl. Acad. Sci. 2017, 114 (9), E1698–E1706. 10.1073/pnas.1615771114.

(7) Chen, H.-Y.; Weng, I.-C.; Hong, M.-H.; Liu, F.-T. Galectins as Bacterial Sensors in the Host Innate Response. Curr. Opin. Microbiol. 2014, 17, 75–81. 10.1016/j.mib.2013.11.006.

(8) Grazier, J. J.; Sylvester, P. W. Role of Galectins in Metastatic Breast Cancer. In Breast Cancer; Mayrovitz, H. N., Ed.; Exon Publications: Brisbane (AU), 2022.

(9) Ahmed, R.; Anam, K.; Ahmed, H. Development of Galectin-3 Targeting Drugs for Therapeutic Applications in Various Diseases. Int. J. Mol. Sci. 2023, 24 (9), 8116. 10.3390/ijms24098116.

(10) Argüeso, P.; Guzman-Aranguez, A.; Mantelli, F.; Cao, Z.; Ricciuto, J.; Panjwani, N. Association of Cell Surface Mucins with Galectin-3 Contributes to the Ocular Surface Epithelial Barrier. J. Biol. Chem. 2009, 284 (34), 23037–23045. 10.1074/jbc.M109.033332.

(11) Argüeso, P.; Mauris, J.; Uchino, Y. Galectin-3 as a Regulator of the Epithelial Junction: Implications to Wound Repair and Cancer. Tissue Barriers 2015, 3 (3), 1–6. 10.1080/21688370.2015.1026505.

(12) Uchino, Y.; Mauris, J.; Woodward, A. M.; Dieckow, J.; Amparo, F.; Dana, R.; Mantelli, F.; Argüeso, P. Alteration of Galectin-3 in Tears of Patients with Dry Eye Disease. Am. J. Ophthalmol. 2015, 159 (6), 1027–1035.e3. 10.1016/j.ajo.2015.02.008.

(13) Vijmasi, T.; Chen, F. Y. T.; Balasubbu, S.; Gallup, M.; McKown, R. L.; Laurie, G. W.; McNamara, N. A. Topical Administration of Lacritin Is a Novel Therapy for Aqueous-Deficient Dry Eye Disease. Invest. Ophthalmol. Vis. Sci. 2014, 55 (8), 5401–5409. 10.1167/iovs.14-13924.

(14) Pflugfelder, S. C.; De Paiva, C. S.; Li, D. Q.; Stern, M. E. Epithelial-Immune Cell Interaction in Dry Eye. Cornea 2008, 27 (SUPPL. 1), 1–7. 10.1097/ICO.0b013e31817f4075.

(15) Craig, J. P.; Nichols, K. K.; Akpek, E. K.; Caffery, B.; Dua, H. S.; Joo, C. K.; Liu, Z.; Nelson, J. D.; Nichols, J. J.; Tsubota, K.; Stapleton, F. TFOS DEWS II Definition and Classification Report. Ocul. Surf. 2017, 15 (3), 276–283. 10.1016/j.jtos.2017.05.008.

(16) Ochieng, J.; Leite-Browning, M. L.; Warfield, P. Regulation of Cellular Adhesion to Extracellular Matrix Proteins by Galectin-3. Biochem. Biophys. Res. Commun. 1998, 246 (3), 788–791. 10.1006/bbrc.1998.8708.

(17) Fukumori, T.; Takenaka, Y.; Yoshii, T.; Kim, H.-R. C.; Hogan, V.; Inohara, H.; Kagawa, S.; Raz, A. CD29 and CD7 Mediate Galectin-3-Induced Type II T-Cell Apoptosis. Cancer Res. 2003, 63 (23), 8302–8311.

(18) Feuk-Lagerstedt, E.; Jordan, E. T.; Leffler, H.; Dahlgren, C.; Karlsson, A. Identification of CD66a and CD66b as the Major Galectin-3 Receptor Candidates in Human Neutrophils1. J. Immunol. 1999, 163 (10), 5592–5598. 10.4049/jimmunol.163.10.5592.

(19) Movassaghi, C. S.; Sun, J.; Jiang, Y.; Turner, N.; Chang, V.; Chung, N.; Chen, R. J.; Browne, E. N.; Lin, C.; Schweppe, D. K.; Malaker, S. A.; Meyer, J. G. Recent Advances in Mass Spectrometry-Based Bottom-Up Proteomics. Anal. Chem. 2025. 10.1021/acs.analchem.4c06750.

(20) Bagdonaite, I.; Malaker, S. A.; Polasky, D. A.; Riley, N. M.; Schjoldager, K.; Vakhrushev, S. Y.; Halim, A.; Aoki-Kinoshita, K. F.; Nesvizhskii, A. I.; Bertozzi, C. R.; Wandall, H. H.; Parker, B. L.; Thaysen-Andersen, M.; Scott, N. E. Glycoproteomics. Nat. Rev. Methods Primer 2022, 2 (1). 10.1038/s43586-022-00128-4.

(21) Rangel-Angarita, V.; Malaker, S. A. Mucinomics as the Next Frontier of Mass Spectrometry. ACS Chem. Biol. 2021, 16 (10), 1866–1883. 10.1021/acschembio.1c00384.

(22) Cioce, A.; Calle, B.; Rizou, T.; Lowery, S. C.; Bridgeman, V. L.; Mahoney, K. E.; Marchesi, A.; Bineva-Todd, G.; Flynn, H.; Li, Z.; Tastan, O. Y.; Roustan, C.; Soro-Barrio, P.; Rafiee, M. R.; Garza-Garcia, A.; Antonopoulos, A.; Wood, T. M.; Keenan, T.; Both, P.; Huang, K.; Parmeggian, F.; Snijders, A. P.; Skehel, M.; Kjær, S.; Fascione, M. A.; Bertozzi, C. R.; Haslam, S. M.; Flitsch, S. L.; Malaker, S. A.; Malanchi, I.; Schumann, B. Cell-Specific Bioorthogonal Tagging of Glycoproteins. Nat. Commun. 2022, 13 (1). 10.1038/s41467-022-33854-0.

(23) Shon, D. J.; Malaker, S. A.; Pedram, K.; Yang, E.; Krishnan, V.; Dorigo, O.; Bertozzi, C. R. An Enzymatic Toolkit for Selective Proteolysis, Detection, and Visualization of Mucin-Domain Glycoproteins. Proc. Natl. Acad. Sci. U. S. A. 2020, 117 (35), 21299–21307. 10.1073/pnas.2012196117.

(24) Shane M. Finn; Keira E. Mahoney; Taryn M. Lucas; Valentina Rangel-Angarita; Ryan J. Chen; Stacy A. Malaker. GlycoFASP: A Universal Method to Prepare Complex Mixtures for O-Glycoproteomic Analysis. ChemRxiv 2025. 10.7868/80424857017030112.

(25) Riley, N. M.; Malaker, S. A.; Driessen, M. D.; Bertozzi, C. R. Optimal Dissociation Methods Differ for N- A Nd O-Glycopeptides. J. Proteome Res. 2020, 19 (8), 3286–3301. 10.1021/acs.jproteome.0c00218.

(26) Sanghi, S.; Kumar, R.; Lumsden, A.; Dickinson, D.; Klepeis, V.; Trinkaus-Randall, V.; Frierson, H. F.; Laurie, G. W. cDNA and Genomic Cloning of Lacritin, a Novel Secretion Enhancing Factor from the Human Lacrimal Gland. J. Mol. Biol. 2001, 310 (1), 127–139. 10.1006/jmbi.2001.4748.

(27) Karnati, R.; Laurie, D. E.; Laurie, G. W. Lacritin and the Tear Proteome as Natural Replacement Therapy for Dry Eye. Exp. Eye Res. 2013, 117, 39–52. 10.1016/j.exer.2013.05.020.

(28) Samudre, S.; Lattanzio, F. A.; Lossen, V.; Hosseini, A.; Sheppard, J. D.; McKown, R. L.; Laurie, G. W.; Williams, P. B. Lacritin, a Novel Human Tear Glycoprotein, Promotes Sustained Basal Tearing and Is Well Tolerated. Invest. Ophthalmol. Vis. Sci. 2011, 52 (9), 6265–6270. 10.1167/iovs.10-6220.

(29) Wang, W.; Jashnani, A.; Aluri, S. R.; Gustafson, J. A.; Hsueh, P.-Y.; Yarber, F.; McKown, R. L.; Laurie, G. W.; Hamm-Alvarez, S. F.; MacKay, J. A. A Thermo-Responsive Protein Treatment for Dry Eyes. J. Control. Release Off. J. Control. Release Soc. 2015, 199, 156–167. 10.1016/j.jconrel.2014.11.016.

(30) Sharifian Gh., M.; Norouzi, F.; Sorci, M.; Zaidi, T. S.; Pier, G. B.; Achimovich, A.; Ongwae, G. M.; Liang, B.; Ryan, M.; Lemke, M.; Belfort, G.; Gadjeva, M.; Gahlmann, A.; Pires, M. M.; Venter, H.; Harris, T. E.; Laurie, G. W. Lacritin Cleavage-Potentiated Targeting of Iron - Respiratory Reciprocity Promotes Bacterial Death. J. Biol. Chem. 2025, 301 (5), 108455. 10.1016/j.jbc.2025.108455.

(31) McKown, R. L.; Coleman Frazier, E. V.; Zadrozny, K. K.; Deleault, A. M.; Raab, R. W.; Ryan, D. S.; Sia, R. K.; Lee, J. K.; Laurie, G. W. A Cleavage-Potentiated Fragment of Tear Lacritin Is Bactericidal. J. Biol. Chem. 2014, 289 (32), 22172–22182. 10.1074/jbc.M114.570143.

(32) Wang, J.; Wang, N.; Xie, J.; Walton, S. C.; McKown, R. L.; Raab, R. W.; Ma, P.; Beck, S. L.; Coffman, G. L.; Hussaini, I. M.; Laurie, G. W. Restricted Epithelial Proliferation by Lacritin via PKCα-Dependent NFAT and mTOR Pathways. J. Cell Biol. 2006, 174 (5), 689–700. 10.1083/jcb.200605140.

(33) Wang, W.; Despanie, J.; Shi, P.; Edman, M. C.; Lin, Y.-A.; Cui, H.; Heur, M.; Fini, M. E.; Hamm-Alvarez, S. F.; MacKay, J. A. Lacritin-Mediated Regeneration of the Corneal Epithelia by Protein Polymer Nanoparticles. J. Mater. Chem. B 2014, 2 (46), 8131–8141. 10.1039/C4TB00979G.

(34) Wang, N.; Zimmerman, K.; Raab, R. W.; McKown, R. L.; Hutnik, C. M. L.; Talla, V.; Tyler, M. F.; Lee, J. K.; Laurie, G. W. Lacritin Rescues Stressed Epithelia via Rapid Forkhead Box O3 (FOXO3)-Associated Autophagy That Restores Metabolism*. J. Biol. Chem. 2013, 288 (25), 18146–18161. 10.1074/jbc.M112.436584.

(35) Willcox, M. D. P.; Argüeso, P.; Georgiev, G. A.; Holopainen, J. M.; Laurie, G. W.; Millar, T. J.; Papas, E. B.; Rolland, J. P.; Schmidt, T. A.; Stahl, U.; Suarez, T.; Subbaraman, L. N.; Uçakhan, O. Ö.; Jones, L. TFOS DEWS II Tear Film Report. Ocul. Surf. 2017, 15 (3), 366–403. 10.1016/j.jtos.2017.03.006.

(36) Chang, V.; Mahoney, K. E.; Lian, I.; Chen, R.; Chung, N.; Utheim, T. P.; Karlsson, N. G.; Malaker, S. A. In-Depth Analysis of the Tear Fluid Glycoproteome Reveals Diverse Lacritin Glycosylation and Spliceoforms. J. Biol. Chem. 2025, 301 (9). 10.1016/j.jbc.2025.110580.

(37) Joeh, E.; O’leary, T.; Li, W.; Hawkins, R.; Hung, J. R.; Parker, C. G.; Huang, M. L. Mapping Glycan-Mediated Galectin-3 Interactions by Live Cell Proximity Labeling. 10.1073/pnas.2009206117/-/DCSupplemental.

(38) Quantitative proteomics reveal the anti-tumour mechanism of the carbohydrate recognition domain of Galectin-3 in Hepatocellular carcinoma | Scientific Reports. https://www.nature.com/articles/s41598-017-05419-5 (accessed 2025-10-18).

(39) Obermann, J.; Priglinger, C. S.; Merl-Pham, J.; Geerlof, A.; Priglinger, S.; Götz, M.; Hauck, S. M. Proteome-Wide Identification of Glycosylation-Dependent Interactors of Galectin-1 and Galectin-3 on Mesenchymal Retinal Pigment Epithelial (RPE) Cells§. Mol. Cell. Proteomics 2017, 16 (8), 1528–1546. 10.1074/mcp.M116.066381.

(40) C. Dange, M.; S. Bhonsle, H.; K. Godbole, R.; K. More, S.; M. Bane, S.; J. Kulkarni, M.; D. Kalraiya, R. Mass Spectrometry Based Identification of Galectin-3 Interacting Proteins Potentially Involved in Lung Melanoma Metastasis. Mol. Biosyst. 2017, 13 (11), 2303–2309. 10.1039/C7MB00260B.

(41) Cederfur, C.; Malmström, J.; Nihlberg, K.; Block, M.; Breimer, M. E.; Bjermer, L.; Westergren-Thorsson, G.; Leffler, H. Glycoproteomic Identification of Galectin-3 and −8 Ligands in Bronchoalveolar Lavage of Mild Asthmatics and Healthy Subjects. Biochim. Biophys. Acta BBA - Gen. Subj. 2012, 1820 (9), 1429–1436. 10.1016/j.bbagen.2011.12.016.

(42) Galectin-3 drives glycosphingolipid-dependent biogenesis of clathrin-independent carriers | Nature Cell Biology. https://www.nature.com/articles/ncb2970 (accessed 2025-10-18).

(43) Georgiev, G. A.; Gh, M. S.; Romano, J.; Teixeira, K. L. D.; Struble, C.; Ryan, D. S.; Sia, R. K.; Kitt, J. P.; Harris, J. M.; Hsu, K. L.; Libby, A.; Odrich, M. G.; Suárez, T.; McKown, R. L.; Laurie, G. W. Lacritin Proteoforms Prevent Tear Film Collapse and Maintain Epithelial Homeostasis. J. Biol. Chem. 2021, 296, 1–16. 10.1074/jbc.RA120.015833.

(44) Francisco, V. V.; Romano, J. A.; McKown, R. L.; Green, K.; Zhang, L.; Raab, R. W.; Ryan, D. S.; Hutnik, C. M. L.; Frierson, H. F.; Laurie, G. W. Tissue Transglutaminase Is a Negative Regulator of Monomeric Lacritin Bioactivity. Invest. Ophthalmol. Vis. Sci. 2013, 54 (3), 2123–2132. 10.1167/iovs.12-11488.

(45) Afshari, A. R.; Chang, V.; Thomsson, K. A.; Höglund, J.; Browne, E. N.; Karadzhov, G.; Mahoney, K. E.; Lucas, T. M.; Rangel-Angarita, V.; Ryberg, H.; Gidwani, K.; Pettersson, K.; Rolfson, O.; Björkman, L. I.; Eisler, T.; Schmidt, T. A.; Jay, G. D.; Malaker, S. A.; Karlsson, N. G. Glycoproteoforms of Osteoarthritis-Associated Lubricin in Plasma and Synovial Fluid. Mol. Cell. Proteomics 2025, 24 (3), 100923. 10.1016/j.mcpro.2025.100923.

(46) Zhou, L.; Beuerman, R. W.; Choi, M. C.; Shao, Z. Z.; Xiao, R. L.; Yang, H.; Tong, L.; Liu, S.; Stern, M. E.; Tan, D. Identification of Tear Fluid Biomarkers in Dry Eye Syndrome Using iTRAQ Quantitative Proteomics. J. Proteome Res. 2009, 8 (11), 4889–4905. 10.1021/pr900686s.

(47) Green-Church, K. B.; Nichols, K. K.; Kleinholz, N. M.; Zhang, L.; Nichols, J. J. Investigation of the Human Tear Film Proteome Using Multiple Proteomic Approaches. Mol. Vis. 2008, 14 (March), 456–470.

(48) Mahoney, K. E.; Chang, V.; Lucas, T. M.; Maruszko, K.; Malaker, S. A. Mass Spectrometry-Compatible Elution Technique Enables an Improved Mucin-Selective Enrichment Strategy to Probe the Mucinome. Anal. Chem. 2024, 96 (13), 5242–5250. 10.1021/acs.analchem.3c05762.

(49) Wan, S. J.; Ma, S.; Evans, D. J.; Fleiszig, S. M. J. Resistance of the Murine Cornea to Bacterial Colonization during Experimental Dry Eye. PLOS ONE 2020, 15 (5), e0234013. 10.1371/journal.pone.0234013.

(50) Halim, A.; Westerlind, U.; Pett, C.; Schorlemer, M.; Rüetschi, U.; Brinkmalm, G.; Sihlbom, C.; Lengqvist, J.; Larson, G.; Nilsson, J. Assignment of Saccharide Identities through Analysis of Oxonium Ion Fragmentation Profiles in LC-MS/MS of Glycopeptides. J. Proteome Res. 2014, 13 (12), 6024–6032. 10.1021/pr500898r.

(51) Pirro, M.; Mohammed, Y.; de Ru, A. H.; Janssen, G. M. C.; Tjokrodirijo, R. T. N.; Madunić, K.; Wuhrer, M.; van Veelen, P. A.; Hensbergen, P. J. Oxonium Ion Guided Analysis of Quantitative Proteomics Data Reveals Site-Specific o-Glycosylation of Anterior Gradient Protein 2 (Agr2). Int. J. Mol. Sci. 2021, 22 (10). 10.3390/ijms22105369.

(52) Characterization of the in vivo forms of lacrimal-specific proline-rich proteins in human tear fluid - Fung - 2004 - PROTEOMICS - Wiley Online Library. https://analyticalsciencejournals.onlinelibrary.wiley.com/doi/10.1002/pmic.200300849 (accessed 2025-10-18).

(53) Kavanaugh, D.; Kane, M.; Joshi, L.; Hickey, R. M. Detection of Galectin-3 Interaction with Commensal Bacteria. Appl. Environ. Microbiol. 2013, 79 (11), 3507–3510. 10.1128/AEM.03694-12.

(54) Nielsen, J. S.; McNagny, K. M. Novel Functions of the CD34 Family. J. Cell Sci. 2008, 121 (22), 3683–3692. 10.1242/jcs.037507.

(55) Liao, C.; Wang, Q.; An, J.; Chen, J.; Li, X.; Long, Q.; Xiao, L.; Guan, X.; Liu, J. CD44 Glycosylation as a Therapeutic Target in Oncology. Front. Oncol. 2022, 12, 883831. 10.3389/fonc.2022.883831.

(56) Dimitroff, C. J.; Lee, J. Y.; Fuhlbrigge, R. C.; Sackstein, R. A Distinct Glycoform of CD44 Is an L-Selectin Ligand on Human Hematopoietic Cells. Proc. Natl. Acad. Sci. 2000, 97 (25), 13841–13846. 10.1073/pnas.250484797.

(57) Clark, M. C.; Baum, L. G. T Cells Modulate Glycans on CD43 and CD45 during Development and Activation, Signal Regulation, and Survival. Ann. N. Y. Acad. Sci. 2012, 1253 (1), 58–67. 10.1111/j.1749-6632.2011.06304.x.

(58) Ng, W.; Loh, A. X. W.; Teixeira, A. S.; Pereira, S. P.; Swallow, D. M. Genetic Regulation of MUC1 Alternative Splicing in Human Tissues. Br. J. Cancer 2008, 99 (6), 978–985. 10.1038/sj.bjc.6604617.

(59) Moniaux, N.; Escande, F.; Batra, S. K.; Porchet, N.; Laine, A.; Aubert, J.-P. Alternative Splicing Generates a Family of Putative Secreted and Membrane-Associated MUC4 Mucins. Eur. J. Biochem. 2000, 267 (14), 4536–4544. 10.1046/j.1432-1327.2000.01504.x.

(60) Ma, P.; Beck, S. L.; Raab, R. W.; McKown, R. L.; Coffman, G. L.; Utani, A.; Chirico, W. J.; Rapraeger, A. C.; Laurie, G. W. Heparanase Deglycanation of Syndecan-1 Is Required for Binding of the Epithelial-Restricted Prosecretory Mitogen Lacritin. J. Cell Biol. 2006, 174 (7), 1097–1106. 10.1083/jcb.200511134.

(61) Dias-Teixeira, K.; Horton, X.; McKown, R.; Romano, J.; Laurie, G. W. The Lacritin-Syndecan-1-Heparanase Axis in Dry Eye Disease. In Advances in Experimental Medicine and Biology; Springer, 2020; Vol. 1221, pp 747–757. 10.1007/978-3-030-34521-1_31.

(62) Zhang, Y.; Wang, N.; Raab, R. W.; McKown, R. L.; Irwin, J. A.; Kwon, I.; Van Kuppevelt, T. H.; Laurie, G. W. Targeting of Heparanase-Modified Syndecan-1 by Prosecretory Mitogen Lacritin Requires Conserved Core GAGAL plus Heparan and Chondroitin Sulfate as a Novel Hybrid Binding Site That Enhances Selectivity. J. Biol. Chem. 2013, 288 (17), 12090–12101. 10.1074/jbc.M112.422717.

(63) Kawashima, M.; Kawakita, T.; Inaba, T.; Okada, N.; Ito, M.; Shimmura, S.; Watanabe, M.; Shinmura, K.; Tsubota, K. Dietary Lactoferrin Alleviates Age-Related Lacrimal Gland Dysfunction in Mice. PLOS ONE 2012, 7 (3), e33148. 10.1371/journal.pone.0033148.

(64) Flanagan, J. L.; Willcox, M. D. P. Role of Lactoferrin in the Tear Film. Biochimie 2009, 91 (1), 35–43. 10.1016/j.biochi.2008.07.007.

(65) Karav, S.; German, J. B.; Rouquié, C.; Le Parc, A.; Barile, D. Studying Lactoferrin N-Glycosylation. Int. J. Mol. Sci. 2017, 18 (4), 1–14. 10.3390/ijms18040870.

(66) Zlatina, K.; Galuska, S. P. The N-Glycans of Lactoferrin: More than Just a Sweet Decoration. Biochem. Cell Biol. 2021, 99 (1), 117–127. 10.1139/bcb-2020-0106.

(67) Kijlstra, A.; Kuizenga, A.; Van Der Velde, M.; Van Haeringen, N. J. Gel Electrophoresis of Human Tears Reveals Various Forms of Tear Lactoferrin. Curr. Eye Res. 1989, 8 (6), 581–588. 10.3109/02713688908995757.

(68) Baksheeva, V. E.; Tiulina, V. V.; Iomdina, E. N.; Petrov, S. Yu.; Filippova, O. M.; Kushnarevich, N. Yu.; Suleiman, E. A.; Eyraud, R.; Devred, F.; Serebryakova, M. V.; Shebardina, N. G.; Chistyakov, D. V.; Senin, I. I.; Mitkevich, V. A.; Tsvetkov, P. O.; Zernii, E. Yu. Tear nanoDSF Denaturation Profile Is Predictive of Glaucoma. Int. J. Mol. Sci. 2023, 24 (8), 7132. 10.3390/ijms24087132.

(69) Schmelter, C.; Brueck, A.; Perumal, N.; Qu, S.; Pfeiffer, N.; Grus, F. H. Lectin-Based Affinity Enrichment and Characterization of N-Glycoproteins from Human Tear Film by Mass Spectrometry. Molecules 2023, 28 (2), 648. 10.3390/molecules28020648.

(70) Cao, X.; Ren, Y.; Lu, Q.; Wang, K.; Wu, Y.; Wang, Y.; Zhang, Y.; Cui, X.; Yang, Z.; Chen, Z. Lactoferrin: A Glycoprotein That Plays an Active Role in Human Health. Front. Nutr. 2023, 9, 1018336. 10.3389/fnut.2022.1018336.

(71) Kell, D. B.; Heyden, E. L.; Pretorius, E. The Biology of Lactoferrin, an Iron-Binding Protein That Can Help Defend Against Viruses and Bacteria. Front. Immunol. 2020, 11. 10.3389/fimmu.2020.01221.

(72) Justis, B. M.; Coburn, C. E.; Tyler, E. M.; Showalter, R. S.; Dissler, B. J.; Li, M.; McNamara, N. A.; Laurie, G. W.; McKown, R. L. Development of a Quantitative Immunoassay for Tear Lacritin Proteoforms. Transl. Vis. Sci. Technol. 2020, 9 (9), 1–12. 10.1167/tvst.9.9.13.

