## Supplemental Information for "Mapping galectin-3 ligands in tear fluid establishes spliceoform-dependent lacritin binding"

### Supplementary Table

Supplementary Table 1: Glycoproteomic analysis of Gal-3 enriched tear fluid and Gal-3 lectin blot densitometry data

### Supplementary Figures

Supplementary Figure 1. Gal-3 affinity enrichment MS workflow
Supplementary Figure 2. Individual patient MS chromatograms and oxonium ion analysis
Supplementary Figure 3. Annotated MS2 spectrum of lacritin glycopeptide
Supplementary Figure 4. Extracted ion chromatograms for lacritin isoforms across all three patients

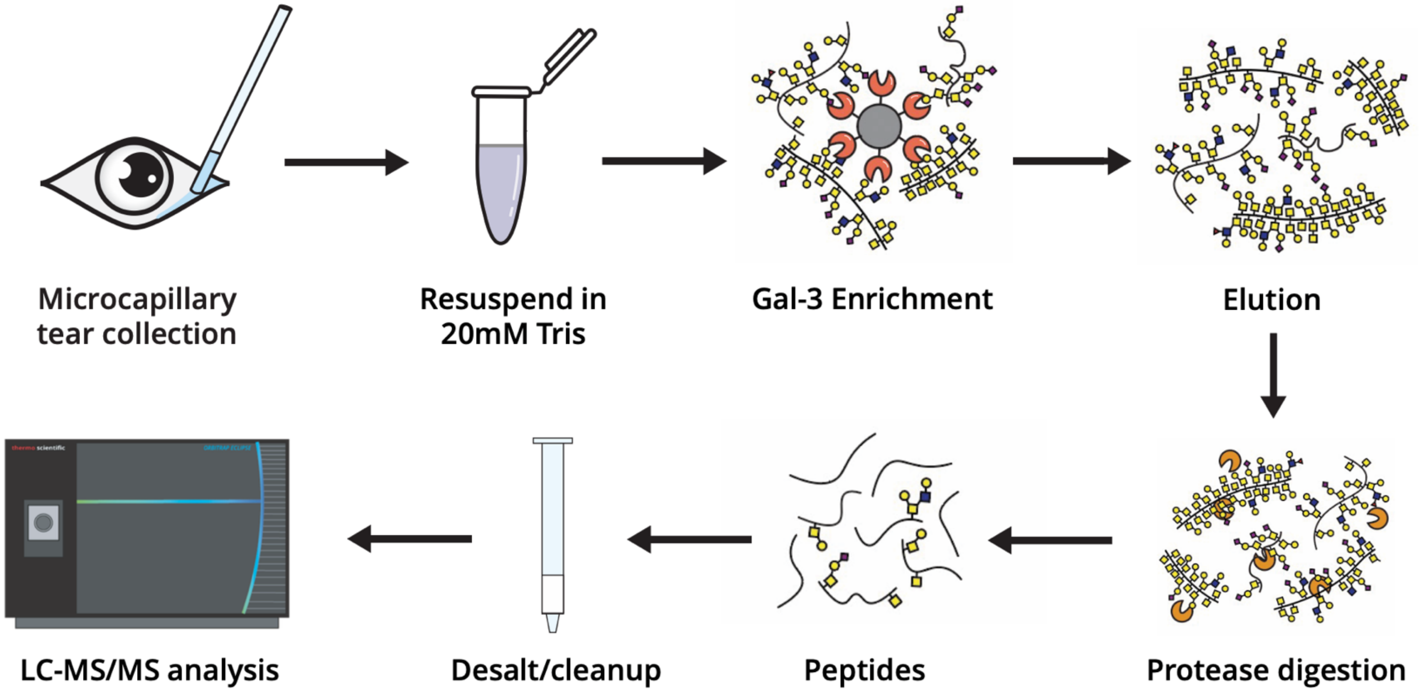


**Supplementary Figure 1. Gal-3 affinity enrichment MS workflow**
Tears from three healthy patients were collected by microcapillary tubes and processed separately from each other. Briefly, 4 μL of tear fluid is resuspended in 20mM Tris (pH=8) and then incubated with Gal-3 conjugated to NHS beads. Post-enrichment, glycoproteins and proteins are digested with trypsin and mucinase SmE before LC-MS/MS analysis with an orbitrap eclipse.

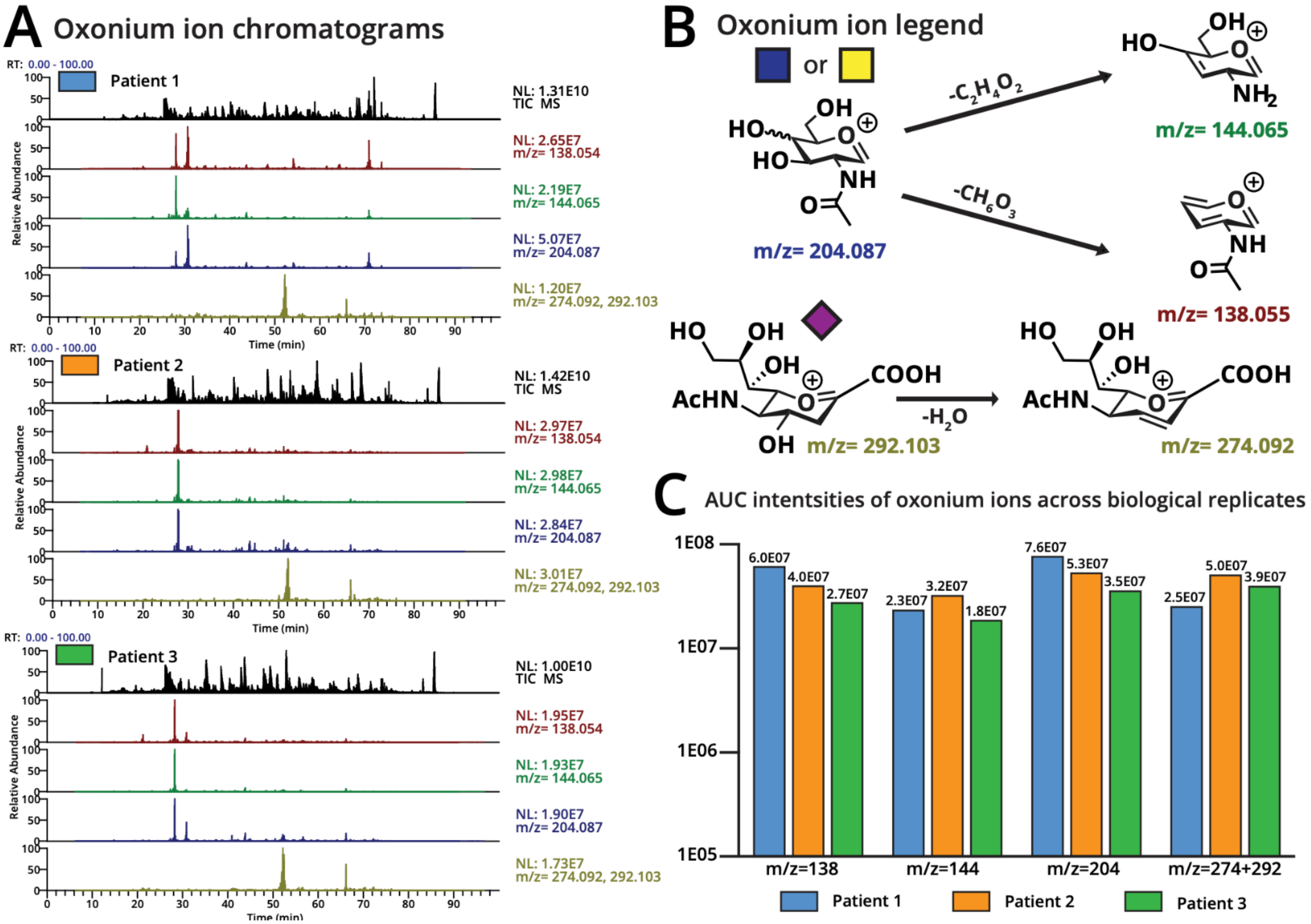


**Supplementary Figure 2. Individual patient MS chromatograms and oxonium ion analysis**
(**A**) Total ion chromarogram and oxonium ion traces for each individual patient across a 90 minute run. (**B**) Oxonium ion legend showing how oxonium ions are formed during higher energy collisional dissociation. (**C**) Area under the curve (AUC) intensities of oxonium ions across each individual patient.

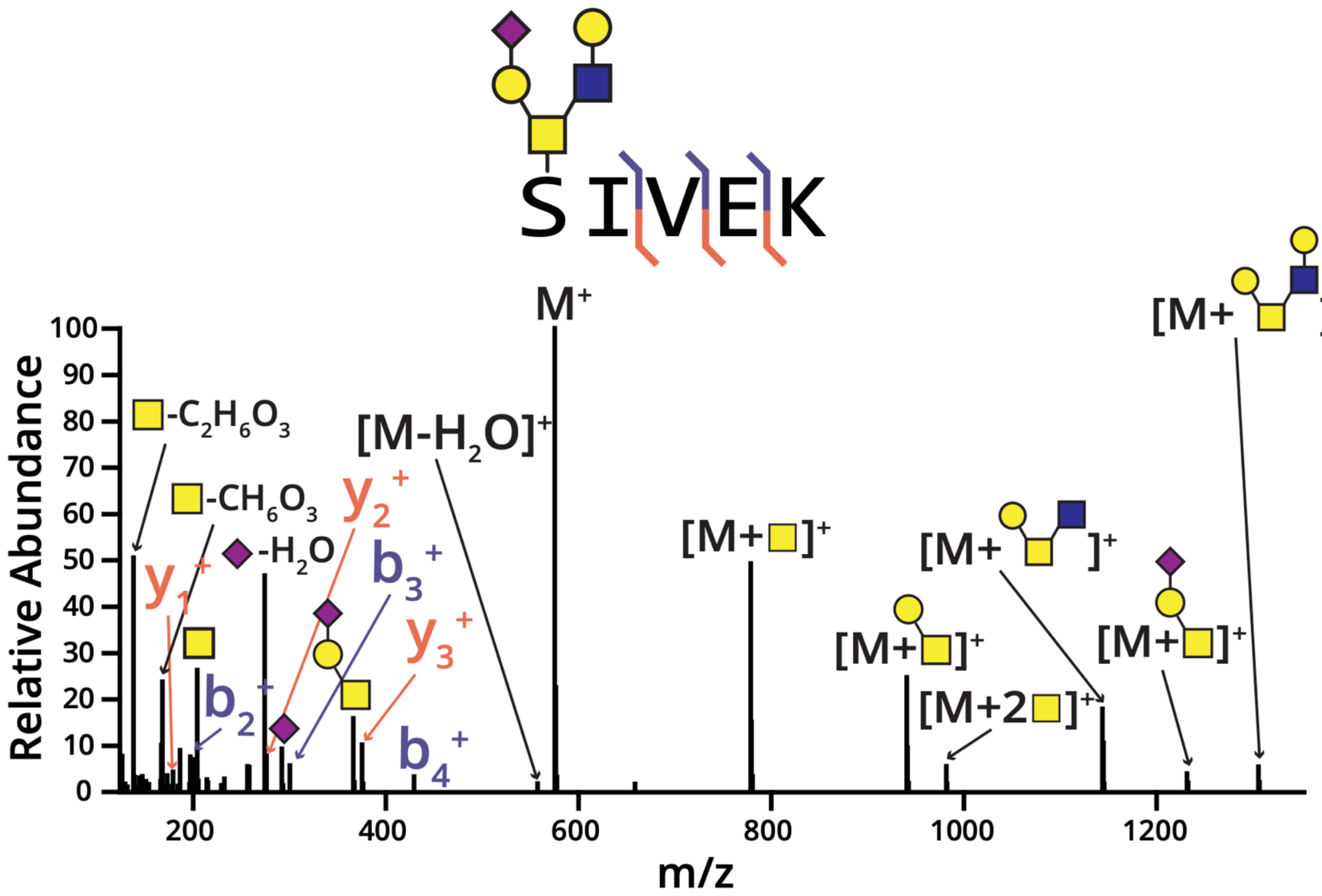


**Supplementary Figure 3. Annotated MS2 spectrum of lacritin glycopeptide**

Lacritin glycopeptide S[H2N2A1] was detected by HCD fragmentation. The MS2 spectrum resulting from HCD shows b-ions (blue) and y-ions (orange) to support the peptide sequence assignment and glycan composition.

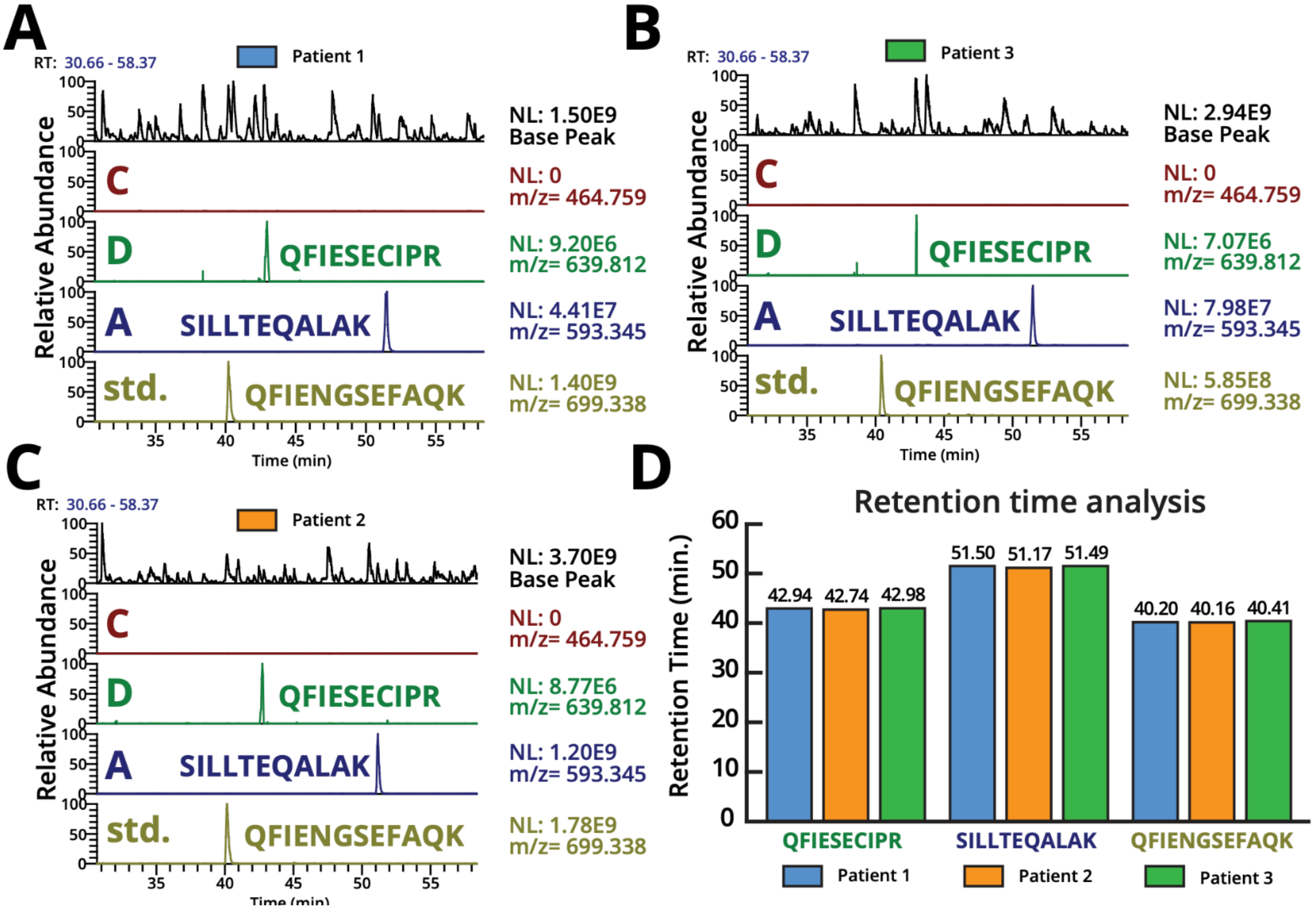


**Supplementary Figure 4. Extracted ion chromatograms (XICs) for lacritin isoforms across all three patients**

Gal-3-enriched tear fluid was subjected to digestion with SmE mucinase and trypsin followed by MS analysis and manual data validation. **(A-C)** Extracted ion chromatogram and the base peak chromatogram (top) for the peptides QELNPLSK (m/z 464.759, z=2), QFIESCIPR (m/z 639.812 z=2), SILLTEQALAK (m/z=593.345, z=2), and QFIENGSEFAQK (m/z=699.338, z=2). These peptides correspond to isoform-specific tryptic peptides for spliceoform C,D, A and the peptide labeled std. was found in all runs and used to normalize AUC intensities for comparison of peptide intensities across runs. (**D**) Retention time analysis of each detected peptide is shown across all three patients, highlighting low variability in retention time.
